# Dynamic constriction and fission of ER membranes by reticulon

**DOI:** 10.1101/472829

**Authors:** Javier Espadas, Diana Pendin, Rebeca Bocanegra, Artur Escalada, Giulia Misticoni, Tatiana Trevisan, Ariana Velasco del Olmo, Sergio Bova, Borja Ibarra, Peter I. Kuzmin, Anna V. Shnyrova, Vadim A. Frolov, Andrea Daga

## Abstract

The endoplasmic reticulum (ER) is a continuous cell-wide membrane network. Network formation has been widely associated with proteins producing membrane curvature and fusion, such as reticulons and atlastin. Regulated network fragmentation, occurring in different physiological contexts, is less understood. We found that the ER network has an embedded fragmentation mechanism based upon the ability of reticulons to produce fission of elongating network branches. In *Drosophila*, fission is counterbalanced by atlastin-driven fusion, with their imbalance leading to ER fragmentation. Live imaging of ER network dynamics upon ectopic expression of *Drosophila* reticulon linked fission to augmented membrane friction. Consistently, *in vitro* analysis revealed that purified reticulon produced velocity-dependent constriction and fission of lipid nanotubes pulled from a flat reservoir membrane. Fission occurred at elongation rates and pulling force ranges intrinsic to the ER network, thus suggesting a novel principle of organelle morphology regulation where the dynamic balance between fusion and fission is governed by membrane motility.

---

The endoplasmic reticulum (ER) comprises two uninterrupted domains, the nuclear envelope and the peripheral ER. The peripheral ER is composed by structural elements with different membrane curvature and topology, from flat sheets and reticulated tubules to complex fenestrated structures. These elements are distributed throughout the cytoplasm of the eukaryotic cell as a membrane network enclosing a single lumen^1–3^. Network maintenance requires homotypic membrane fusion mediated by the atlastin-family of dynamin-related GTPases^4,5^. Suppression of atlastin fusogenic activity leads to ER fragmentation^4^ thus revealing an endogenous mechanism aimed at the reduction of ER connectedness. The existence of this mechanism has been confirmed by several reports showing ER disassembly during mitosis^6–8^, reversible fragmentation of the ER both in neurons^9^ and other cell types^10,11^ and fragmentation of the ER prior to autophagic degradation^12,13^. While no dedicated molecular machinery has been linked to ER fragmentation, few experimental observations suggest an involvement of reticulons^12–14^, highly conserved integral ER membrane proteins implicated in shaping and stabilizing the tubular ER^15–18^. Notably, mutations in both reticulon and atlastin, the major players so far implicated in shaping of the ER network, have been linked to the neurodegenerative disorder hereditary spastic paraplegia^19,20^, corroborating their participation in coordinated functional and pathological pathways.

Overexpression of members of the Yop1 and reticulon families of proteins has been reported to cause severe constriction of ER branches^15,21^ and ER fragmentation^14^. Fragmentation could proceed via the breakage of ER tubules, implicating high local curvature stress and membrane fission, and/or via the shedding of small vesicles^14^, a process whose significance in ER fragmentation, however, is not understood. Tubule fission would naturally antagonize the fusogenic activity of atlastin in the ER, making fusion/fission balance a paradigm in intracellular organelle maintenance. Despite the broad association between reticulons and ER fragmentation, direct involvement of reticulons has not been shown and the mechanism(s) of fragmentation remains obscure. Furthermore, reticulon-induced curvatures have been mechanistically linked to construction, not fragmentation of the tubular ER network both *in vitro* and *in vivo*^5,15,22^. In agreement with a role in supporting formation rather than fragmentation of the tubular ER, purified reticulons reconstituted into lipid vesicles induced membrane curvatures insufficient to produce membrane fission^15,23^.

Here, we reveal the mechanism underlying reticulon membrane activity that unifies these seemingly contradictory observations. We found that in vivo *Drosophila* reticulon (Rtnl1) is responsible for ER fragmentation both at endogenous amounts, when unchallenged by the absence of atlastin, and upon overexpression by promoting ER fission. Corroborating *in vivo* results, purified Rtnl1 reconstituted into dynamic lipid nanotubes created curvatures ranging from moderate, as reported earlier^15^, to those causing spontaneous membrane fission. *In vivo* this ability of Rtnl1 to induce membrane fission is counterbalanced by atlastin, with the interplay between these proteins exerting the core control on total curvature and connectedness of the ER network in a living organism.

To address the function of reticulon in vivo we used *Drosophila* because, unlike vertebrates, it has a single reticulon gene (*Rtnl1*) and examined its genetic interaction with atlastin (*atl*). Homozygous *Rtnl1^1^* null flies^24^ are viable and normal while homozygous *atl^2^* null individuals ^25^ die at the pupa stage with a 2% rate of escapers. Surprisingly, we found that combining these two null mutations in homozygosity resulted in 84% adult survival (Fig. 1a). Hence, removal of *Rtnl1* substantially alleviates the lethality associated with depletion of *atl*, indicating that a robust antagonistic genetic interaction between atlastin and reticulon exists in *Drosophila*. This interaction was confirmed in the fly eye where RNAi-mediated loss of Rtnl1 in an eye expressing wild type atlastin resulted in increased severity of the atlastin-dependent small eye phenotype (Extended Data, Fig. 1a) and in the nervous system where the lethality produced by D42-Gal4 driven overexpression of atlastin in motor neurons was markedly enhanced in the *Rtnl1^1^* mutant background. EM tomography-based 3D reconstruction of the ER network in *atl^2^* neurons showed disconnected ER elements (Fig. 1b, Movie 1,2), supporting earlier observations of ER fragmentation after loss of atlastin^4^. Remarkably, depletion of Rtnl1 in the *atl^2^* null background restored a normal ER structure: ER network organization in *Rtnl1^1^/atl^2^* neurons resembles closely that of control neurons comprising interconnected tubular and sheet-like elements (Fig. 1b, Movies 3-6). The observation that removal of Rtnl1 in the *atl^2^* null background restores both viability and ER shape strongly indicates that Rtnl1 is the force driving the morphological alterations and fragmentation of the ER caused by loss of the fusogenic activity of atlastin. Importantly, our data demonstrate that balancing the activities of atlastin and Rtnl1 is critical not only for the maintenance of the ER architecture but also for organism survival.

**Figure 1.**
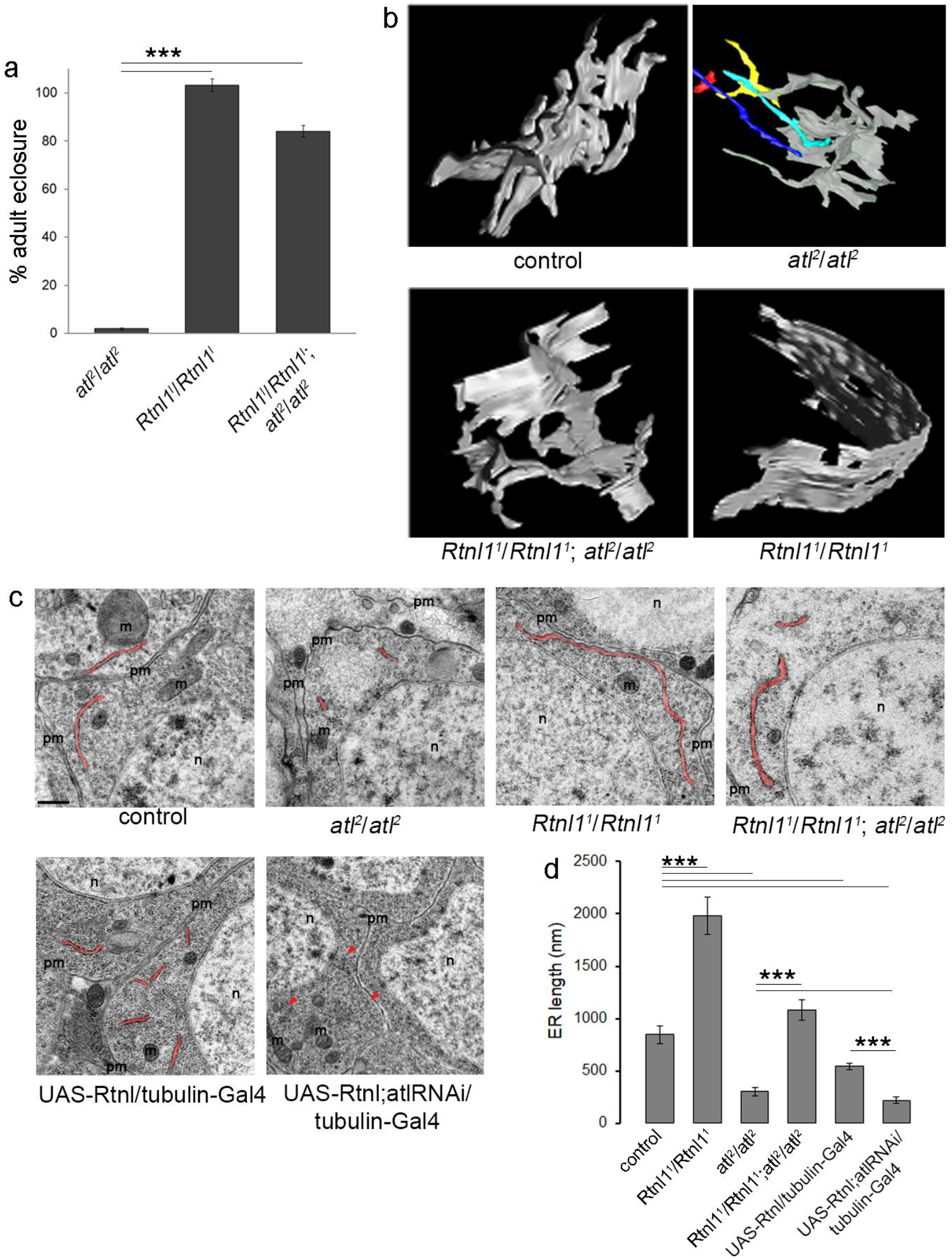
The genetic antagonism between Rtnl1 and atlastin in *Drosophila* is reflected in morphological alterations of the ER. **a,** The histogram displays the percentage of surviving adults, expressed as the ratio of observed over expected individuals, for the indicated genotypes. **b,** EM tomography-based 3D reconstruction of portions of ER network from neurons of the indicated genotypes. ER elements not connected are shown in color. Scale bar 200 nm. **c,** Representative EM images of ventral ganglion neuronal bodies of the indicated genotypes highlighting ER profiles in red. Scale bar 0.5 μm. pm, plasma membrane; m, mitochondria; n, nucleus. **d,** Average length of ER profiles measured on thin EM sections shown in (**c**).

The effects of reticulons on ER morphology have been linked to their ability to bend membranes: reticulons convert flat ER sheets into more curved tubular structures^16,26^ thereby increasing the total curvature of ER membranes. In agreement with this notion, EM tomography in *Rtnl1^1^* mutant neurons revealed elongated unbranched ER sheets (Fig. 1b, Movie 7,8). This effect can be quantified using a simple metrics: the average length of ER profiles, corresponding to a cut through sheet-like structures, on thin EM sections (Fig. 1c). When compared with controls, *Rtnl1^1^* mutant neurons displayed elongated ER profiles (1980±177nm vs 846±85 nm), as previously reported for a different cell type^27^. Notably, ER profile elongation in *Rtnl1^1^* flies was suppressed by re-expression of transgenic Rtnl1 (Extended Data, Fig. 1b,c). Coherently, Rtnl1 overexpression in a wild type background led to profile length reduction (542±28nm) (Fig. 1c, d). EM analyses revealed that the effect of Rtnl1 on profile length is normally compensated by the activity of atlastin. In fact, ER profile length was substantially shortened in neurons lacking atlastin (303±40nm)^4^ thus demonstrating that loss of atlastin or Rtnl1 change profile length in opposite direction. Consistent with this observation, profile length reduction by Rtnl1 overexpression was exacerbated by simultaneous downregulation of atlastin. Indeed, UAS-atl-RNAi,UAS-Rtnl1/tub-Gal4 neurons showed a significant decrease of average ER profile length (to 217±31nm) when compared to UAS-Rtnl1/tub-Gal4, where endogenous atlastin can actively oppose Rtnl1 function, as well as to UAS-atl-RNAi/tub-Gal4 alone (Fig 1c, d) where profile length reduction is due to uncontested endogenous Rtnl1. Even more striking than profile length decrease was the paucity of ER observed in UAS-atl-RNAi,UAS-Rtnl1/tub-Gal4 neurons (Fig 1c), indicating that much of the network was broken up in small, unidentifiable fragments thus making our quantitative analysis biased towards visible, longer profiles. Finally, ER profile length in *Rtnl1^1^/atl^2^* double mutant neurons was comparable to that of control neurons (1081±99nm) demonstrating reciprocal compensation of the mutant phenotypes (Fig. 1c, d) and thus confirming the opposing effect of the two genes. These results demonstrate that atlastin offsets the reduction of ER profile length mediated by endogenous or transgenic Rtnl1, making this parameter a measure of the functional balance between Rtnl1 and atl *in vivo*.

Both atlastin and reticulons were demonstrated to create membrane curvature^18^, so their antagonistic effect on the ER profile length cannot be explained by the diminished local membrane bending activity. Hence, we analyzed the functional interaction between atlastin and Rtnl1 at the network level. STED fluorescence microscopy of whole larva brain showed that in neurons both atlastin downregulation and Rtnl1 overexpression caused relocation of the luminal ER marker GFP-KDEL to bright punctae in the perinuclear region (Fig. 2a). Accumulation of the luminal marker in these punctae was evident from the analysis of the fluorescence intensity distribution over the cytoplasm (Fig. 2b). Similar bright structures emerged in larva muscles both upon atlastin downregulation and overexpression of Rtnl1 (Fig. 2c). We showed earlier by fluorescence loss in photobleaching (FLIP) that appearance of these punctae correlated with fragmentation of the ER lumen, since in *atl^2^* muscles free diffusion of GFP-KDEL in the ER is abolished^4^. FLIP analysis of muscles ectopically co-expressing Rtnl1 and GFP-KDEL revealed a comparable loss of the diffusional exchange of GFP-KDEL between bleached and non-bleached ER regions (Fig. 2d). Therefore, when Rtnl1 overpowers atlastin, either by its overexpression or by loss of atlastin, the ER lumen becomes disconnected and broken into distinct fragments. Fragmentation puts a natural limit on the size of continuous ER elements thus providing a plausible explanation for the diminished length of the ER profiles upon shifting Atl/Rtnl1 balance towards the latter (Fig. 1d). Hence, the profile length reflects not only morphology (i.e. tubulation) but also topology changes, with atlastin promoting fusion and Rtnl1 fragmentation of ER membranes.

**Figure 2.**
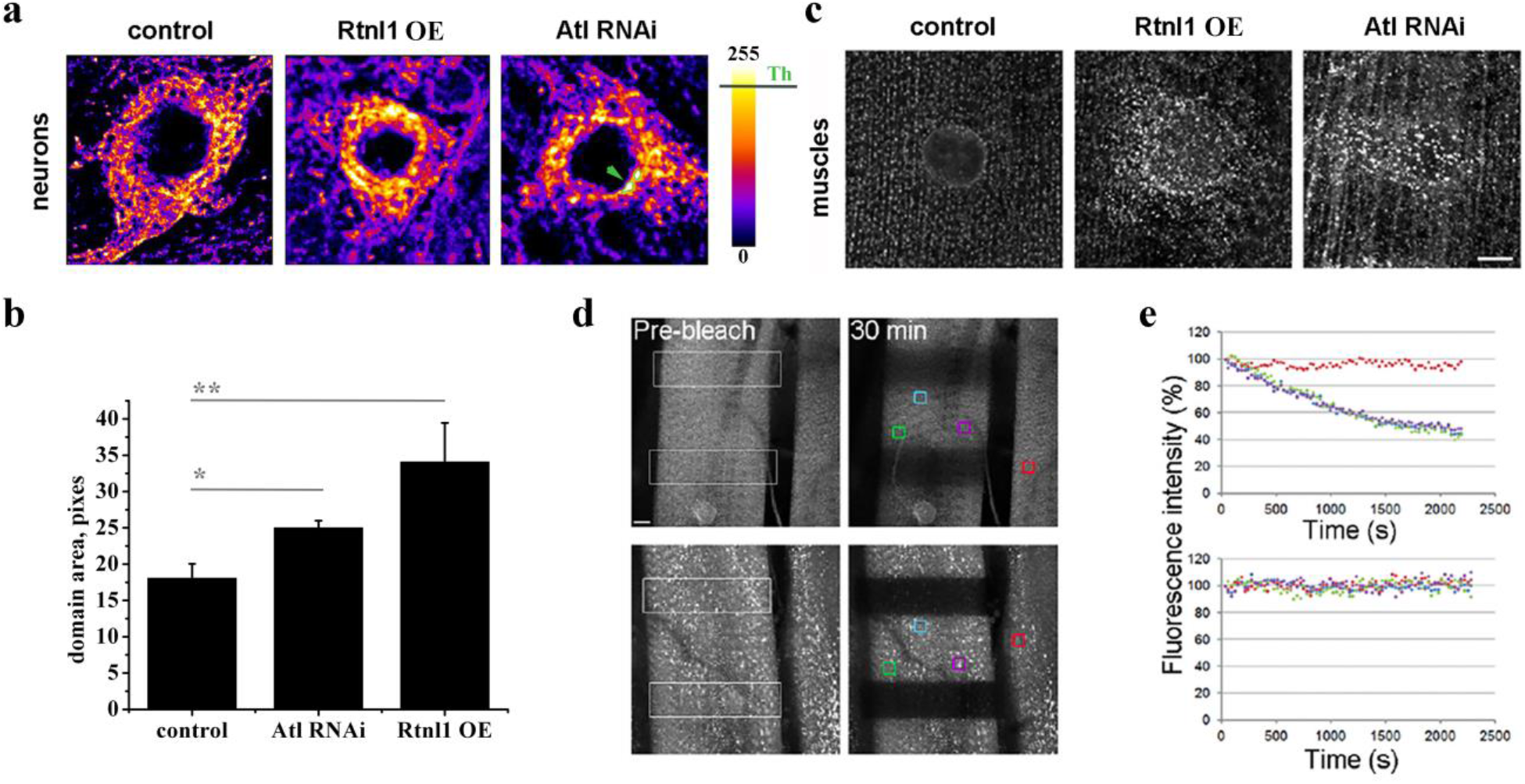
Rtnl1 overexpression and loss of atlastin give rise to comparable defects in ER distribution and connectedness in *Drosophila*. **a,** Deconvolved confocal STED projections showing comparable changes in the ER network appearance produced by Rtnl1 overexpression (Rtnl1 OE) and knockdown of atlastin (atl-RNAi) in neurons labeled with the ER marker GFP-KDEL (upper row). The pseudo-color representation highlights emergence of bright fluorescent domains (examples marked by the arrow) in Rtnl1 overexpression and atl-RNAi. Scale bar 5 μm. **b,** Rtnl1 overexpression and atl-RNAi show increase of the total area of bright fluorescent domains (calculated at the brightness threshold Th=200). **c,** Emergence of the bright fluorescent punctae in third instar larva muscle labeled by GFP-KDEL in Rtnl1 overexpression and atl-RNAi. Scale bar 10 μm. **d,** Representative images of FLIP performed by repetitive photobleaching of two regions (white outline box) in control and Rtnl1 overexpressing muscles labeled with GFP-KDEL (left). Scale bar 10 μm. **e,** Rates of fluorescence loss in four independent regions of the muscle (color boxes) were quantified and graphed. The red box was chosen on an adjacent unbleached muscle as a control.

As the above results show that ER morphology changes are associated with the amount of Rtnl1 unopposed by Atl, we resorted to ectopic expression of Rtnl1 in COS-7 cells whose expanse tubular ER network enables direct assessment of fragmentation. As in the fly, overexpression of Rtnl1 in COS-7 cells caused transformation of the continuous ER network into bright punctae (Fig. 3a). Both the lumen and the membranes of these fluorescent domains were visually unconnected (Fig. 3b). Remarkably, live imaging of the transforming network at early stages of transfection revealed scissions of individual ER tubules both near the ends and in the middle portion of the tubules (Fig. 3c, red arrow, Movies 9, 10), pointing to membrane fission as the mechanism underlying ER fragmentation. We scored a disconnection event as fission when two disconnected parts of the tubule snapped in opposite directions, indicating involvement of substantial axial force (Fig. 3c; Movies 9, 10). Such forces, created by molecular motors pulling on the ER network^28^, are intrinsic to actively remodeling, dynamic regions of the ER network, such as the peripheral ER where Rtnl1 driven fragmentation is the most apparent (Fig. 3a). Interestingly, Rtnl1 overexpression also caused significant slowing of the retraction of disconnected ER branches (Fig. 3d). Although impaired retraction is consistent with stabilization of tubular ER branches by reticulons, it may also indicate the presence of a frictional barrier impairing nanotube pulling, an effect previously linked to facilitation of membrane fission^29^.

**Figure 3.**
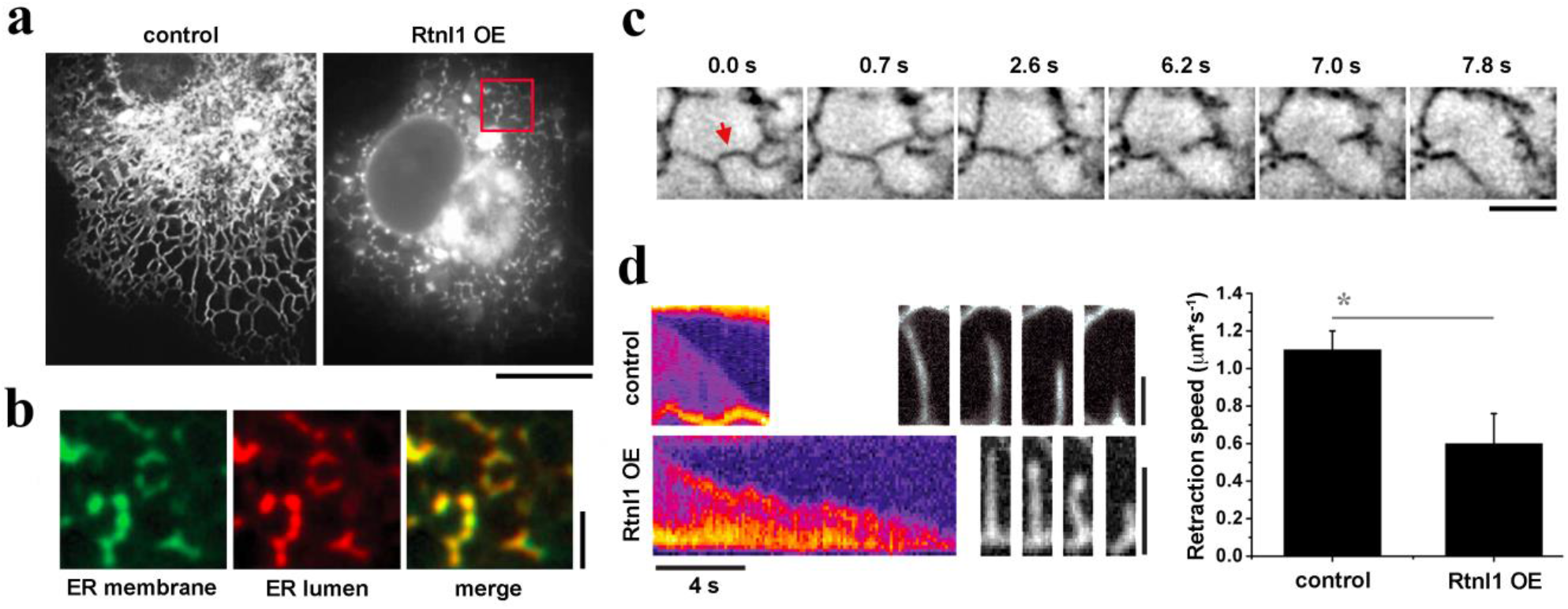
Altered dynamics and fission of ER branches during Rtnl1-driven fragmentation of tubular ER network in COS-7 cells. **a,** Retraction and constriction of the ER network (labeled by mCHERRY-KDEL in COS-7 cells expressing Rtnl1. Global ER constriction results in the appearance of multiple bright punctae of mCHERRY-KDEL fluorescence. Scale bar 10 μm. **b,** Blow-ups of the ER in COS-7 cells co-transfected with Rtnl1-HA, mCHERRY-KDEL and Rtnl1-GFP (24h post-transection). Both fluorescence markers localize to visibly disconnected punctae. **c,** Image sequence showing scission (red arrow) of an ER branch in a Rtnl1-expressing COS-7 cell. **d,** Kymographs with corresponding selected frame sequences showing retraction of ER branches in control (upper row, 1s between frames) and Rtnl1-expressing (lower row, 2s between frames) cells. The bar-graph on the right shows branch retraction speed (the error bars show SD, control n=10, Rtnl1 n=5).

To explore coupling between membrane dynamics and fission, we reconstituted purified Rtnl1 into lipid nanotubes mimicking dynamic ER branches. We pulled the tubes from proteo-lipid bilayers formed on a surface of silica microbeads by proteo-liposome deposition (Extended Data, Fig. 2). The nanotube appearance monitored by fluorescence microscopy depended upon Rtnl1: lipid ratio. At low ratio (up to 1:300) uniform cylindrical tubes were formed, resembling pure lipid nanotubes. At higher protein:lipid (1:150) bulged and constricted regions appeared during pulling (Fig. 4a). Rtnl1 incorporation into the nanotubes was verified using the labelled protein (Fig. 4a, lower panel). Measurements of the axial force during pulling with constant speed (V_t_) revealed that constriction is accompanied by a linear force growth followed by a plateau (Fig. 4b), a behaviour characteristic of viscous drag^29–32^. Consistent with observations in COS-7 experiments, the axial force increase caused fission of the nanotube (Fig. 4b, c, Movie 11). During the limited elongations tested in our experiments (ΔL<10μm) only a fraction of the tubes broke. Decreasing the length of non-broken tubes at a constant V_t_ caused the axial force to temporarily drop to zero as indicate by nanotube sagging (Fig. 4d, upper panel), an effect reminiscent of the slowed retraction of ER branches in COS-7 cells overexpressing Rtnl1 (Fig. 3d).

**Figure 4.**
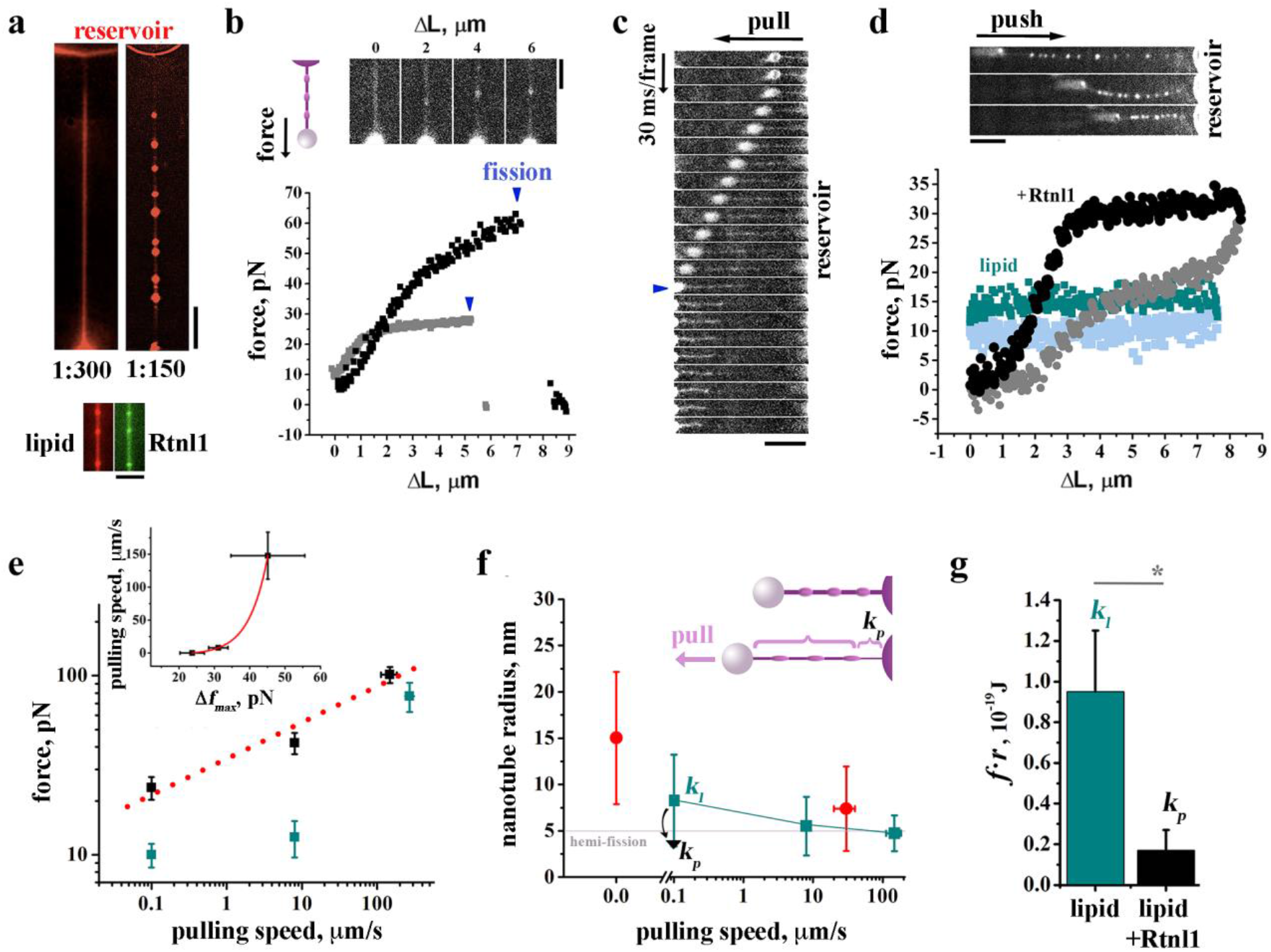
Constriction-by-friction mechanism of Rtnl1-driven membrane fission. **a,** Representative images of lipid nanotubes (Rh-DOPE fluorescence) obtained at 1:300 (left) and 1:150 (right) Rtnl1/lipid (mol/mol), Scale bar 10 μm. The lower panel shows incorporation of the ALexa488-Rtnl1 (green) into the nanotube (red) at 1:150 protein/lipid. Scale bar 2μm. **b,** Increase of the axial force during elongation of Rtnl1-nanotubes at 0.1 μm/s (grey) and 8 μm/s (black) speeds caused fission seen as an abrupt decrease of the force to zero (arrows). The image sequence shows the nanotube constriction during the elongation (see Extended Data, Fig. 3f). Scale bar 2μm. **c,** Frame sequence (30ms/frame) demonstrating scission (arrow) of Rtnl1-nanotube. Scale bar 5μm. **d,** Changes of the axial force during consecutive elongation/shortening (ΔL) of pure lipid (dark cyan/light blue) and Rtnl1-(black/grey) nanotubes at constant V_t_=8μm/s. The image sequence shows sagging of Rtnl1-constricted membrane nanotube during shortening, Scale bar 5μm. **e,** Dependence of the axial force measured at the moment of fission of Rtnl1-containing tubes (*f*_Rtnl1_, black, red dotted line shows linear regression) or maximal force measured during 10μm elongation of lipid tubes (*f*_lip_, cyan) on the elongation speed (V_t_). The insert shows the force difference Δ*f_max_* = (*f*_*Rtnl*1_ – *f_lip_*) dependence on V_t_, where the red line is the logarithmic fit (Supplementary Information). The error bars show SE. **f,** Radial constriction of the membrane nanotube measured by fluorescence microscopy (red, the error bars show SD, n=10) and recalculated from the force increase (shown in **e**) using either *k_l_* (cyan) or *k_p_* (black). **g,** Effective bending rigidity calculated as 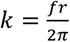 for Rtnl1 (black, *k_p_*) and pure lipid (cyan, *k_l_*) nanotubes; *f* and *r* values are from Extended Data Fig. 3e and 4b.

The extension-contraction cycle revealed a hysteresis in force-length dependence, large in Rtnl1-containing tubes and virtually absent in pure lipid nanotubes (Fig. 4d). This behaviour is consistent with the viscous drag, considerably increased by the presence of Rtnl1, opposing both retraction and lengthening of the nanotube. The increase of the axial force during pulling measured at the point of fission (Fig. 4b) showed characteristically weak dependence on V_t_^29,32^ (Fig. 4e, black). For physiologically relevant elongation speeds the force ranged from 23.7±3.4 pN at V_t_ =0.1 μm/s to 41.5±5.4 pN V_t_ =8μm/s, comparable with the forces (20-40pN) reported in nanotubes pulled from the ER and Golgi membrane networks^33^. Hence *in vitro* Rtnl1 probabilistically causes nanotube scission at elongation speeds and forces existent within the ER network. At V_t_ =0.1 μm/s 15 out of 32 tubes broke 36.4±2.8s after the beginning of elongation (Fig. 4b, grey curve), corresponding to an average 3.6 μm extension of the tube prior to fission. Due to the limited elongation length, at V_t_=8μm/s the nanotubes were subjected to elevated stress for much shorter time (0.64±0.49s) than they were at 0.1 μm/s speed and only 5 out 17 nanotubes broke (Fig. 4b, black curve). This indicates stochastic coupling between membrane stresses and fission^34^. At higher, non-physiological V_t_ (above 100μm/s) 12 out of 12 Rtnl1-containing nanotubes broke as the force reached 108±18pN (Fig. 4e). Such high-speed pulling produced similar force increase but, markedly, no scission (in 21 out of 21 cases) of pure lipid nanotubes (Fig. 4e, cyan). To directly compare the force effect, we subjected lipid nanotubes to elevated forces (above 30pN) for the same amount of time (0.74±0.23s) as Rtnl1 nanotubes elongating at 8μm/s. None of 15 lipid tubes broke as compared with 30% fission rate in the Rtnl1 experiments (see above). Hence, a moderate force increase per se is insufficient to trigger fission of a lipid bilayer tube^23^. It has been suggested that protein presence in the nanotube could facilitate formation of a pore even under moderate tensile stress, thus leading to fission via membrane poration^29^. However, we found that in Rtnl1-containing tubes force-driven constriction is enhanced by the intrinsic curvature activity of Rtnl1 and/or its oligomers^15,35^. In static nanotubes (V_t_=0), Rtnl1-induced curvature stabilized nanotube constriction at 15.0±2.7nm radius (Fig. 4f, red, Extended Data, Fig. 3e), a value comparable with earlier *in vitro* observations^15^. During elongation, the pulling force acted on such pre-constricted parts of the nanotube bringing their curvature close to the hemi-fission threshold (Fig. 4f, red)^23^ and thus enabling fission along an alternative, pore-free path^36,37^

The reduction of the radius (*R_t_*) of pre-constricted regions is linked to the pulling force increase as 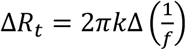, where *k* is the effective bending rigidity modulus of the nanotube membrane ^29,38^; Supplementary Information). ΔR_t_ calculated with *k* of pure lipid nanotubes (*k_l_*, Fig. 4g) and the fission force values (Fig. 4e, black), matched that directly measured by fluorescent microscopy (Fig 4f, cyan and red, Supplementary Information). Hence, the portions of nanotube pre-constricted by Rtnl1 retained lipid elasticity, indicating sparse Rtnl1 coverage^15^. We emphasize that such coverage enables dynamic transformations of the nanotube, in striking contrast with rigid protein scaffolds, e.g. Dynamin1 helix completely encaging lipid nanotubes^39^. Notably, Dynamin1 produced static constriction similar to Rtnl1, yet, Dynamin1 scaffolds retain their geometry during elongation thereby preventing force-driven constriction and ensuing fission (Extended Data, Fig. 3f). Limited Rtnl1 coverage implies moderate friction resistance (Supplementary Information), enabling nanotube pulling at physiologically relevant elongation speeds and forces. Increasing Rtnl1:lipid ratio to 1:80 completely impaired nanotube production (12 out of 12 cases). Thus, the ability to combine dynamic and static constrictions in the context of a living ER network is specific for Rtnl1 and is likely related to its unique molecular architecture comprising curvature- and friction-creating elements.

Interaction between the two constriction modes is more complex near the reservoir membrane, where elongating nanotubes appeared visibly thinner (Fig 4c). Our theoretical analysis linked the effect to curvature-driven sorting of Rtnl1 between the reservoir membrane and the emergent part of the nanotube near the reservoir (Fig. 4f, cartoon, Supplementary Information)^40^. The sorting, seen as force relaxation upon step elongation of the tube (Extended Data Fig. 4a, b), caused 6-fold reduction of *k* compared to stationary nanotubes (*k*_p_, Fig 4g), thus greatly facilitating the constriction (Fig. 4f, black arrow, Supplementary Information). In the ER, Rtnl1 sorting would facilitate local constriction during short step elongation (e.g. Fig. 3c) and enhance fission of the tubules pulled from low curvature parts of the network, such as sheets, explaining their prevalence at the late stages of Rtnl1 over-expression (Fig. 3a). Furthermore, the sorting also explained the weak growth of the fission force with V_t_ (Fig. 4e) by shear thinning ^38^ (Supplementary Information). Subtracting the lipid contribution to the force increase (linear with V_t_ at high speeds^31^ (Fig. 4e, cyan) revealed that *f*(V_t_) is logarithmic at high speeds (Fig. 4e, insert, Supplementary Information). By fitting *f*(V_t_) we found that Rtnl1 caused ~100-fold increase of the viscosity al low physiologically relevant speeds. The large membrane-inserting reticulon-homology domain, fully spanning the outer and partially the inner lipid leaflets, is likely to produce such a viscous drag, further enhanced by reticulon oligomerization^41^.

We conclude that Rtnl1 utilizes two synergistic modes of membrane curvature, static and dynamic, whose combination can produce either stable membrane curvature or fission, depending on intrinsic membrane dynamics. Our findings imply that in a living cell, the ability of Rtnl1 to produce membrane constriction is inseparable from its ability to produce fission. Under these circumstances, atlastin-mediated fusion becomes necessary to maintain overall connectedness in a dynamic ER network. The intrinsic sensitivity to membrane movement suggests a novel paradigm of dynamic regulation of ER topology, linking membrane fusion and fission with membrane motility. This paradigm implies that ER fragmentation, a process crucial in physiological conditions, for example maintenance of ER morphology and ER-phagy, and likely involved in neuropathological processes^42^ can be implicitly controlled by multiple factors connected to ER motility and stresses, with Rtnl1 constituting the core element of the ER-specific membrane fission machinery.

## Methods

### *Drosophila* genetics and behavioral analysis

Fly culture and transgenesis were performed using standard procedures. Several transgenic lines for UAS-HA-Rtnl1 were generated and tested. *Drosophila* strains used: GMR-Gal4, D42-Gal4, tubulin-Gal4, armadillo-Gal4; pUASp:Lys-GFP-KDEL. UAS-Rtnl1-RNAi lines were obtained from Vienna *Drosophila* RNAi Center (v7866 and v33919). Lifespan experiments were performed with 200 animals for each genotype. Flies were collected 1 day after eclosion and placed in vials containing 50 animals. The animals were maintained at 25°C, transferred to fresh medium every day, and the number of dead flies was counted. Lifespan experiments were repeated 3 times.

### Fluorescence Loss In Photobleaching (FLIP)

FLIP experiments were performed as described^4^. The experiments were repeated at least three times.

### Electron microscopy

*Drosophila* brains were fixed in 4% paraformaldehyde and 2% glutaraldehyde and embedded as described earlier^4^. EM images were acquired from thin sections under a FEI Tecnai-12 electron microscope. EM images of individual neurons for the measurement of the length of ER profiles were collected from three brains for each genotype. At least 20 neurons were analyzed for each genotype. Quantitative analyses were performed with ImageJ software^43^.

### Electron Tomography

Epon-embedded *Drosophila* larva brains were cut transversally to the ventral nerve cord with Leica Ultracut UCT ultramicrotome. 200-250 nm thick serial sections were collected on formvar carbon-coated slot grids and 10 nm colloidal gold particles were deposited on both surfaces for fiducials. Samples were imaged on a FEI Tecnai G2 20 operating at 200 kV (Lab6) with a FEI eagle 2k CCD camera at a nominal magnification of 14500 that resulted in a resolution of 1.5 nm per pixel. FEI single tilt tomography holder was tilted over a range of +/- 65 degree according to a Saxton’s scheme (2 degrees starting angle, for a total of 87 images collected) using the FEI Xplore3d acquisition software. Tilted images alignment, tomography reconstruction (WBP) and tomograms joining was done with the IMOD software package^44^. Endoplasmic reticulum structures were rendered by manually segmenting the membranes of ER profiles using IMOD software^45^.

### Super resolution imaging

Third instar larva brain neurons were imaged in fixed larvae preparations. Stacks of optical sections (300 nm apart) from the neurons were obtained in a Leica TCS STED CW SP8 super-resolution microscope with a 63x 1,40NA oil (n=1,518) objective using an Argon laser with an excitation line at 458 nm, adjusting the pinhole at one Airy unit and depletion laser at 592 nm. Super resolution images were deconvolved with PSF Generator and DeconvolutionLab plug-ins for ImageJ (Biomedical Imaging Group, École Polytechnique Féderale de Laussanne, bigwww.epfl.ch). The Born & Wolf 3D Optical Model generated a theoretical PSF using PSF Generator. Deconvolution was performed with the Richardson-Lucy algorithm using 70-80 iterations. Background subtraction was done before the deconvolution process. In each case, PSF stacks were generated with the same number of z-planes as of the image stacks deconvolved.

### Calculation of the bright puncta area in the super resolution images

The super-resolution images of larva ventral ganglion neurons from control, Rntl1 OE and Atl-RNAi flies were thresholded at 90% of maximal intensity (as shown in Fig. 3a). The resulting binary images were used as the masks defining the bright puncta in the images. The area and mean fluorescence intensity of the puncta were further analyzed using Analyze Particles algorithm of ImageJ^43^.

### Cell culture

COS-7 cells were cultured in DMEM (HyClone, high glucose, from Thermo Scientific) supplemented with 10% fetal bovine serum and 50 μg/ml Gentamicin. For the fluorescence microscopy experiments cells were plated in 35 mm low wall μ-Dishes (Ibidi GmbH) and further transfected with vectors for expression of DsRed-KDEL (Clontech) and Rtnl1-HA (2 μg DNA each) using lipofectamine 2000 (Invitrogen) according to the manufacturer procedure.

### Live imaging

Live-cell imaging was performed using a microscope stage top incubator INUG2 (Tokai Hit) equipped with objective heater to maintain the cells at 37°C with 5% CO_2_. Revolution-DSD confocal system (Andor, Ireland) equipped with 100x 1.45NA objective lense was used to visualize the membrane morphologies and spatial distribution of labeled Rtnl1 in COS-7 cells transfected with DsRed-KDEL and eGFP-Rtnl1.

### Protein expression and purification

The Reticulon1-PB isoform was obtained from the *Drosophila* Genomic Resource Center (LD14068). The Rtnl1-PB cDNA was subcloned into the pGEX-GST-SUMO1 vector. The Rtnl1 protein was expressed in bacteria and affinity purified. Rtnl1 was eluted from the GST-affinity beads by digestion with GST-SENP2 protease overnight at 4°C. Wild-type human Dynamin 1 was produced in Sf9 insect cells and purified as described earlier^34^. Protein concentration was determined by using the BCA assay kit (ThermoFischer Scientific, USA).

### Protein labeling

Purified Rtnl1 was labeled with Alexa Fluor^™^ 488 maleimide (ThermoFischer Scientific, USA) according to the manufacturer’s instructions. Dye excess was removed by using dye removal columns (ThermoFischer Scientific, USA). The labeling efficiency was assayed by absorption measurements and was ~0.2 dye/protein. Dyn1-Atto488 (generously provided by Dr. Sandra Schmid, UTSouthwestern) was used in some of the experiments.

### Large Unilamellar Vesicles (LUVs) preparation

Dioleoyl-phosphatidyl-choline (DOPC), Dioleoyl-phosphatidyl-ethanolamine (DOPE), Dioleoyl-phosphatidyl-serine (DOPS), Rhodamine-DOPE (Rh-DOPE), cholesterol (chol) and phosphatidylinositol 4,5-bisphosphae (PI4,5P_2_), all from Avanti Polar Lipids, were used to prepare the LUVs. For Rtln1 reconstitution we used DOPC:DOPE:DOPS:Chol:Rh-DOPE in 59.5:20:10:10:0,5 mol%, variation of DOPC/DOPE from 3:1 to 1:3 caused no effect on the Rtnl1-curvature and fission activities. In experiments with Dyn1, the lipid composition was DOPE:DOPC:Chol:DOPS:Rh-DOPE:PI4,5P_2_ 39,5:38:10:10:0,5:2 mol%, The lipid stocks mixed in chloroform were dried under a stream of N_2_ gas followed by further drying under vacuum for 120 min. The lipid films were resuspended in working buffer (20 mM HEPES pH 7.4, 150 mM KCl, 1 mM EDTA) to a final total lipid concentration of about 10 mM. For Dyn1 experiments, the film was resuspended in 1 mM Hepes. In both cases, large unilamellar vesicles (LUVs) were formed by 10 freeze-thaw cycles followed by extrusion through polycarbonate filters with 100 nm pore size (Avanti Polar Lipids, USA).

### Rtnl1 reconstitution into LUVs

Preformed LUVs were diluted to 1 g/L (1,6 mM) with working buffer and titrated with Triton X100 to measure the optical density at 540nm and find the optimum for LUV destabilization^46^. The detergent-destabilized liposomes (final concentration ~0,2 mM) were then mixed with Alexa488-Rtnl1 (11μM in working buffer with 10% glycerol). After 15 min of co-incubation at RT with gentle agitation, the detergent was removed with BioBeads^®^ SM-2 adsorbant (BioRad) added four times to the proteo-lipid mixture in the period of 12h as described earlier ^47^. The sample was then centrifuged for 1 hour at 15000xg to remove the BioBeads^®^ and non-incorporated protein. Rtnl1 incorporation into LUVs was measured by SDS-PAGE of the supernatant. Three independently prepared Rtnl1 batches have been used.

### Formation of lipid and proteo-lipid reservoir membranes on silica and polystyrene beads

Proteoliposomes where dialyzed against 1 mM Hepes and 1 mM trehalose solution. 10 μl of the freshly dialyzed proteo-liposome or LUVs in 1 mM Hepes solution were mixed with 2 μl of 40 μm silica or 5 μm polystyrene beads (Microspheres-Nanospheres, USA), deposited on a teflon film in small drops and then dried in vacuum for 20 min. A 10 μL plastic pipette tip was cut from the bottom to approximately 2/3 of its original size. The cut tip was used to take 6 μL of 1M TRH solution buffered with 1 mM Hepes. The tip was carefully detached from the micropipette and a small portion of the beads covered by dried proteo-lipid or lipid films were picked up (by means of a fire closed patch glass capillary) and deposited into the TRH solution from the top of the tip. The tip was then carefully introduced into a home-made humidity chamber and subjected to 10-20 minutes incubation at 60°C. Then the beads were transferred to the observation chamber, pretreated with bovine serum albumin and filled with the working buffer. Upon immersion in the working buffer, lipid or proteo-lipid film swelling was followed as spontaneous GUV or proteo-GUV formation. Proteo-GUVs attached to beads are shown in the Fig. S5. An estimation of protein incorporation into the proteo-GUVs is described in Fig. S6. Optionally, by decreasing the membrane reservoir on the beads, the formation of vesicles was suppressed, and hydrated lamellas around the bead were formed instead.

### Pulling membrane nanotubes from reservoir membrane

Glass micropipettes were prepared with the P-1000 micropipette puller (Sutter Instruments, USA). A streptavidin covered polystyrene bead (2 μm in diameter, Microspheres-Nanospheres, USA) was trapped at the tip of the micropipette by suction and used to pull the tube from the lipid or proteolipid film or GUV whose membrane was previously doped with 0.2% of biotinylated-DOPS lipid (Avanti Lipids, USA). A micro-positioning system based on 461xyz stage and high resolution NanoPZ actuators (Newport, USA) were used for micropipette approximation and pulling maneuvers.

### Fluorescence microscopy of the membrane nanotubes

An Olympus IX71 inverted microscope equipped with iXon-EMCCD camera (Andor), BrightLine filter sets for Alexa488 (490/520nm excitation/emission) and Rhodamine (560/590nm excitation/emission), 150x 1.45NA objective lens and custom-made observation chamber was used to monitor the tubules pulled from lipid and proteo-lipid films. Images were acquired using μManager open source software at 30 or 100 fps^48^. Images were further processed using ImageJ for cropping, background removal and brightness/contrast adjustments^49^.

### Quantification of the nanotube radius

Nanotube radii were calculated form fluorescence intensity calibration of the lipid film on a flat surface as described previously^50^. Briefly, flat supported bilayer was used to find the density of the membrane fluorescence signal (ρ_0_), then the nanotube radius was obtained from the total fluorescence per unit length of the nanotube *Fl* using r=*Fl*/2πρ_0_.

### Force measurements with optical tweezers

A counter propagating dual-beam optical tweezers instrument equipped with light-momentum force sensors was used in these experiments, which is capable of measuring force directly^51^. The two lasers are brought to the same focus through opposite microscope objective lenses generating a single optical trap. Protein-lipid nanotubes were generated *in situ* as follows: Pre-hydrated 5 μm polystyrene beads covered with proteolipid lamellas (as described above) were introduced into the experimental chamber containing 20mM of working buffer at 22 ±1°C. One bead was hold in the optical trap and brought into contact with a 2 μm streptavidin-covered bead immobilized by suction in a micropipette tip. The two beads were separated with an initial constant pulling speed of 0.1 μm/s to form a tube. Extension-shortening cycles were performed on individual tubes at different pulling rates (as indicated in the main text). Below 8 μm s^−1^ the trap was displaced linearly at a fixed calibrated speed. For higher velocities, the pipette was displaced by a coarse positioner away from the optical trap, and the pulling rates were calculated off line as the distance change per unit time. Data was collected with high force (<1pN), position (1-10 nm) and temporal (500 Hz) resolutions. A similar procedure was performed with pure lipid films to test the behaviour of protein-free tubes under force.

## Acknowledgements

This work was partially supported by MINECO BFU2015-70552-P and NIH R01GM121725 to V.A.F., by a 5×1000 grant from the Italian Ministry of Health and Telethon GGP11189 to A.D. and by BFU2015-63714-R to B.I.

## Author contributions

**Conceptualization,** A.D., V.A.F., A.V.S.; **Methodology,** A.D., V.A.F., A.V.S., D.P., B.I.; **Investigation,** J.E., D.P., A.E., G.M., T.T., A.V.d.O., R.B., A.V.S.; **Theoretical analysis** V.A.F., P.I.K.; **Writing – Original Draft**, A.D., V.A.F.; **Writing – Review & Editing**, A.D., V.A.F., J.E., A.V.S., D.P.; **Funding Acquisition,** A.D., V.A.F.; **Resources**, A.D., V.A.F., B.I., S.B.; **Supervision,** A.D., V.A.F., A.V.S., B.I.

